# Gut microbiome signatures of risk and prodromal markers of Parkinson’s disease

**DOI:** 10.1101/2019.12.11.872481

**Authors:** Sebastian Heinzel, Velma T. E. Aho, Ulrike Suenkel, Anna-Katharina von Thaler, Claudia Schulte, Christian Deuschle, Lars Paulin, Sari Hantunen, Kathrin Brockmann, Gerhard W. Eschweiler, Walter Maetzler, Daniela Berg, Petri Auvinen, Filip Scheperjans

## Abstract

**Objectives:** Alterations of the gut microbiome in Parkinson’s disease (PD) have been repeatedly demonstrated. However, little is known about whether such alterations precede disease onset and how they may be related to risk and prodromal markers of PD. We investigated associations of these features with gut microbiome composition.

**Methods:** Established risk and prodromal markers of PD as well as factors related to diet/lifestyle, bowel function and medication were studied in relation to bacterial α-/β-diversity, enterotypes, and taxonomic composition in stool samples of 666 elderly TREND study participants.

**Results:** Among risk and prodromal markers, physical inactivity, constipation and age showed associations with α- and β-diversity, and for both measures subthreshold parkinsonism and physical inactivity showed interaction effects. Moreover, male sex, possible REM-sleep behavior disorder (RBD), smoking as well as body-mass-index, antidiabetic and urate-lowering medication were associated with β-diversity. Physical inactivity and constipation severity were increased in individuals with the *Firmicutes*-enriched enterotype. Subthreshold parkinsonism was least frequently observed in individuals with the *Prevotella*-enriched enterotype. Differentially abundant taxa were linked to constipation, physical inactivity, possible RBD, and subthreshold parkinsonism. Substantia nigra hyperechogenicity, olfactory loss, depression, orthostatic hypotension, urinary/erectile dysfunction, PD family history and the overall prodromal PD probability showed no significant microbiome associations.

**Interpretation:** Several risk and prodromal markers of PD are associated with changes in gut microbiome composition. However, the impact of the gut microbiome on PD risk and potential microbiome-dependent subtypes in the prodrome of PD need further investigation based on prospective clinical and (multi)omics data in incident PD cases.

## INTRODUCTION

The presence of gastrointestinal pathological α-synuclein deposits and constipation in prodromal and clinically established Parkinson’s disease (PD) suggests an integral role of the gut-brain-axis for the early pathogenesis of PD^1–3^. The synucleinopathy is hypothesized to ascend via the vagal nerve from peripheral neurons of the gastrointestinal tract to the brain^4^. Moreover, increased intestinal permeability^5^, elevated stool inflammatory cytokines^6^, and colonic wall inflammation^7^ have been shown in PD patients, and may also represent key gastrointestinal processes in prodromal PD. Mice overexpressing α-synuclein show aggravated motor dysfunction when colonized with intestinal microbiota from PD patients^8^. However, the specific role of gut microbiota for PD and the factors modulating such processes along the microbiota-gut-brain-axis are still largely unknown. The most consistently shown PD-related changes of gut microbial composition include an increase in the relative abundances of *Verrucomicrobiaceae* and *Akkermansia* and a decrease in *Prevotellaceae* and *Prevotella*^9,10^. The latter has also been associated with progressive PD motor symptoms over two years^11^ and with REM-sleep behavior disorder (RBD)^12^, a highly specific prodromal marker of PD. Prospective evidence of several PD risk markers (indicating an increased PD risk) and prodromal markers (indicating an already initiated neurodegenerative process) confirm the concept of a prodromal phase preceding clinically established PD by years or even decades^13^. Based on this evidence, it is possible to define specific predictive values for these markers and to calculate individual prodromal PD probabilities based on age and marker profiles^14,15^. Age^16^ and several markers established in the International Parkinson and Movement Disorder Society (MDS) research criteria for prodromal PD^14,15^ have also been associated with gut microbial changes; these include risk markers such as male sex^17^, diabetes^18^, non-smoking and non-use of caffeine^16,17^ and prodromal markers, such as RBD^12^, constipation^16^ and depression^19^. Moreover, physical inactivity, a risk marker for PD^15^ and for most major chronic diseases occurring frequently in old age^20^, has been reported to affect microbial β-diversity in elderly males^21^ as well as various inflammatory and immune processes^22^. However, these studies focused on microbial associations with single or few PD-related markers, or studied very small samples considering the multitude of other markers and confounding factors that may modulate the gut microbiome. For instance, factors related to diet, bowel function and disease/medication often exert effects on microbial composition^16–18^, and may thus bias findings of marker associations. The present study therefore assessed gut microbial diversity, enterotypes and taxonomic composition and investigated their associations with a comprehensive set of PD risk and prodromal markers, the overall prodromal PD probability and a wide range of other potential confounders in a large sample of elderly individuals.

## METHODS

All subjects were participants of the prospective **T**übingen evaluation of **R**isk factors for **E**arly detection of **N**euro**D**egeneration (TREND) study. The cohort has been enriched regarding an increased PD risk by partly recruiting participants based on the presence of olfactory loss, depression and/or possible RBD. Biennial comprehensive assessments in 1202 individuals have been performed over the last 10 years (www.trend-studie.de/english). Stool samples were collected at the third follow-up of the study and associated with markers/factors assessed at the corresponding wave of assessments. Stool was sampled using collection tubes containing a DNA stabilizer (PSP Spin Stool DNA Plus Kit, STRATEC Molecular), provided using postal services and frozen and stored at −80 °C immediately upon arrival. Samples were available from 745 participants. After excluding individuals taking antibiotic medication (n = 47), patients with PD (n = 11) or atypical/secondary parkinsonism (n = 2), incident cases of PD (n = 3), and individuals with missing dietary and medication data (n = 16), data from 666 individuals were included in the analyses.

Risk and prodromal markers as defined by the recently updated MDS research criteria for prodromal PD were investigated.^15^ We assessed the PD risk markers male sex, substantia nigra hyperechogenicity (transcranial ultrasound), non-smoking, positive PD family history (first degree relatives), physical inactivity (measured as hours of sport/week), type 2 diabetes (diagnosis), and occupational solvent and pesticide exposure (self-reported). Prodromal markers comprised olfactory loss (Sniffin’ Sticks, 16 SS), depression (acute or lifetime diagnosis), constipation (Rome-III criteria, sum score)^23^, possible RBD (RBD screening questionnaire), erectile and urinary dysfunction, symptomatic orthostatic hypotension (the latter three assessed using the UMSARS questionnaire as described previously)^24^. Moreover, motor deficits indicating subthreshold parkinsonism were assessed using the MDS-UPDRS-III^25^ (score >6 after excluding action/postural tremor items; scores from 3 to 6 were defined as borderline motor deficit)^14^. Age and comprehensive individual marker profiles were used to calculate prodromal PD probabilities and to diagnose possible (>50% probability) and probable prodromal PD (>80%) according to the MDS research criteria for prodromal PD^14^. Moreover, body-mass-index (BMI), exhaustion from climbing 3 levels of stairs [no, yes, not capable], education level as well as irritable bowel syndrome, bloating, vomiting/diarrhea, gout, 15 diet-related variables (assessing weekly meat, vegetables, dairy, protein, carbohydrate, and alcohol consumption) and 51 variables related to medication were assessed. Overall, 108 variables were investigated for associations with microbial measures. For the full variable list, specific assessment methods, variable definitions and descriptive statistics, see Supporting material. The study was approved by the local ethical committee (Medical Faculty, University of Tübingen; 444/2019BO2). All participants provided written informed consent.

### Stool sample DNA extraction, library preparation, and sequencing

DNA extraction, library preparation and sequencing were performed at the DNA Sequencing and Genomics Laboratory of the Institute of Biotechnology, University of Helsinki. All samples were stored at −80 °C and randomized in a −20 °C room before bulk DNA extraction with the PSP Spin Stool DNA Plus Kit (STRATEC Molecular). One blank was added per extraction batch to assess potential contamination. After extraction, DNA concentrations were measured with NanoDrop ND-1000 (Thermo Scientific). V3-V4 regions of the bacterial 16S ribosomal RNA gene were amplified in Verity 96-well Thermal Cyclers (Thermo Scientific) using a previously described PCR protocol^11^. The amount of template DNA used for the first PCR ranged between 11.25 ng and 1311.26 ng. Each PCR batch included blank samples for assessment of potential contamination. Dual-indexes were used in the second PCR; these had been selected using BARCOSEL^26^ to allow pooling and sequencing of all samples in one pool run on three separate runs on a MiSeq (Illumina; v3 600 cycle kit, forward/reverse read length 328/278 bases). Thus, each sample was sequenced three times among all the other samples, reducing possible batch effects.

### Sequence analysis

The raw sequence data contained 48,782,168 sequence reads (availability: ENA accession number PRJEB32920). We combined the sequence reads from the three sequencing runs, and then trimmed primers and low-quality sequences with cutadapt (v.1.8.3)^27^ (parameters q=30 and m=160). Merging paired reads, alignment to a reference database (SILVA, v.132), chimera removal, taxonomic classification (reference database: RDP, v.16 PDS), and OTU clustering (“cluster.split” approach) were run with mothur (v.1.40.0)^28^, following the Standard Operating Procedure for MiSeq data^29^. Parameters differing from the SOP were maxlength=500 and maxhomop=8 for the first “screen.seqs”, start=2 and end=17012 for the second “screen.seqs”, diffs=4 for “pre.cluster”, cutoff=70 for “classify.seqs”, and keeping archaeal sequences in “remove.lineage”. Additionally, singleton sequences were removed prior to “classify.seqs” using “split.abund” with cutoff=1. After excluding data from blanks and all OTUs with ≤10 sequence reads, the final data set consisted of 25,390,744 sequence reads (34,082 ± 4,785 per sample).

### Microbial measures

All microbiota analyses were performed using genus-level data. α-diversity was estimated with the inverse Simpson index (R package: phyloseq)^30^. This index summarizes richness (number of different taxonomic units) and evenness (abundance distribution of taxonomic units) for each sample. The measure chosen for β-diversity was Bray-Curtis (BC) dissimilarity (R package: vegan). It quantifies the inter-sample compositional dissimilarity based on both presence-absence of taxonomic units and their abundances. Microbial enterotypes^31^ were determined using the algorithm provided at http://enterotypes.org/. All unclassified taxa (genus level) were excluded from alpha diversity, enterotype and differential abundance analyses. For differential abundance, analyses were also performed for the family-level and OTU-level.

### Statistical analyses

Each of the 108 single markers/factors was investigated for an association with the microbial measures. Variables showing a statistical trend (p<.1) were selected for subsequent multifactorial statistical testing. Final multifactorial models comprised all markers/factors showing marginal effects with p<.1. Linear multiple regressions of α-diversity were performed. Associations of markers/factors with β-diversity were tested using PERMANOVAs (single variable: *adonis*, multifactorial: *adonis2*, both commands from the R package vegan). Moreover, α- and β-diversity were tested for interaction effects between physical inactivity and other risk and prodromal markers of PD. Marker/factor differences between enterotypes were tested using Fisher’s exact tests and Kruskal-Wallis tests, and subsequent multifactorial analyses using multinomial logistic regressions with enterotype as dependent variable. For differential abundance analysis we used DESeq2, a method based on generalized linear models with negative binomial distributions (sequence count data). The DESeq2 model included all covariates accounted for in the final PERMANOVA model for β-diversity, and p-values were adjusted for multiple testing using false-discovery-rate (FDR) corrections. Due to the exploratory nature of the study, for other statistical tests effects with an uncorrected p<.05 were considered significant, while effects with .05≤p<.1 are reported as non-significant statistical trends. We used R (v3.5.1) for all analyses and figures. TREND study data were collected and managed using REDCap electronic data capture tools hosted at the University of Tübingen.

## RESULTS

### Descriptive statistics

The TREND study sample (n = 666) had a mean age (± SD, range) of 68.4 ± 6.3 years (53 - 86). Risk and prodromal marker variables showed the following descriptive statistics: male sex (52.7%), substantia nigra hyperechogenicity (21.6%), positive PD family history (14.7%), physical inactivity (no activity (25.7%); <1 hrs/week (30.9%); 1-2 hrs/week (24.2%); 2-4 hrs/week (8.3%); >4 hrs/week (10.4%)), type 2 diabetes (8.3%), occupational solvent exposure (10.8%), occupational pesticide exposure (1.3%), olfactory loss (19.7%), depression (31.5%), constipation (severity sum score: 3.1 ± 3.9, range 0 – 25), possible RBD (12.6%), erectile dysfunction (in males: 21.5%), urinary dysfunction (5.1%), symptomatic orthostatic hypotension (4.7%), and regarding motor functions, no motor deficit (90.5%), borderline motor deficit (6.0%) and subthreshold parkinsonism (3.5%). Based on prodromal PD probabilities (3.8 ± 11.4%, 0.0 - 89.1%), 15 individuals (2.3%) were diagnosed with possible prodromal PD, 5 individuals (0.8%) with probable prodromal PD.

The most abundant bacterial genus in the subjects’ stool samples was *Bacteroides*, followed by *Faecalibacterium, Gemmiger, Roseburia, Prevotella* and *Ruminococcus* (Fig 1).

**FIGURE 1.**
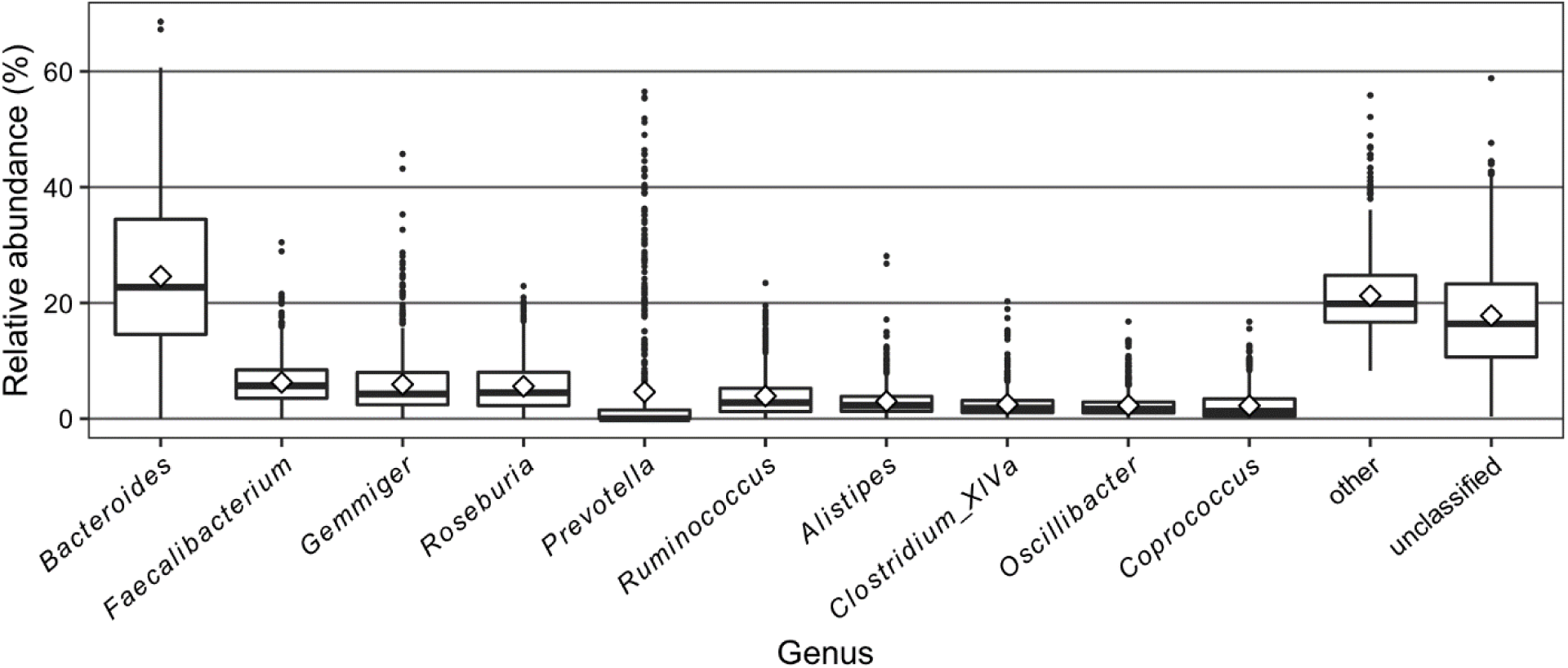
Relative abundances of the 10 most common bacterial genera.

### α-diversity

Alpha diversity on the genus level as indicated by the inverse Simpson index was significantly associated with age, two risk markers, two prodromal markers, and four additional factors (Table 1). Overall, these variables explained 8.0% of α-diversity variance (adjusted R^2^).

**TABLE 1.**
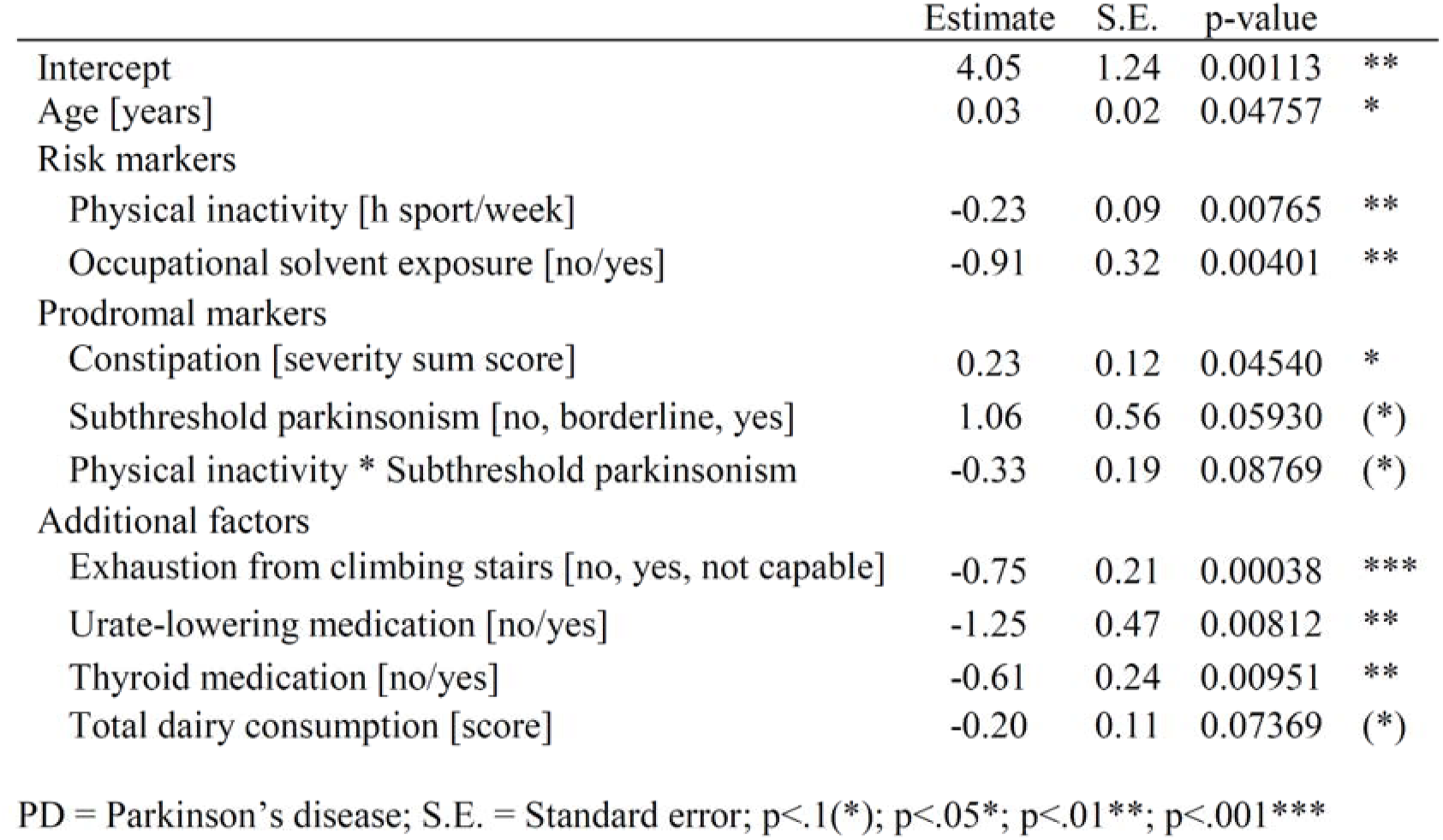
Associations of alpha diversity with risk and prodromal markers of PD and additional factors.

|  | Estimate | S.E. | p-value |  |
| --- | --- | --- | --- | --- |
| Intercept | 4.05 | 1.24 | 0.00113 | ** |
| Age [years] | 0.03 | 0.02 | 0.04757 | * |
| Risk markers |  |  |  |  |
| Physical inactivity [h sport/week] | -0.23 | 0.09 | 0.00765 | ** |
| Occupational solvent exposure [no/yes] | -0.91 | 0.32 | 0.00401 | ** |
| Prodromal markers |  |  |  |  |
| Constipation [severity sum score] | 0.23 | 0.12 | 0.04540 | * |
| Subthreshold parkinsonism [no, borderline, yes] | 1.06 | 0.56 | 0.05930 | (*) |
| Physical inactivity * Subthreshold parkinsonism | -0.33 | 0.19 | 0.08769 | (*) |
| Additional factors |  |  |  |  |
| Exhaustion from climbing stairs [no, yes, not capable] | -0.75 | 0.21 | 0.00038 | *** |
| Urate-lowering medication [no/yes] | -1.25 | 0.47 | 0.00812 | ** |
| Thyroid medication [no/yes] | -0.61 | 0.24 | 0.00951 | ** |
| Total dairy consumption [score] | -0.20 | 0.11 | 0.07369 | (*) |

PD = Parkinson's disease; S.E. = Standard error; $p < .1$ (\*); $p < .05$ \*; $p < .01$ \*\*; $p < .001$ \*\*\*

Increasing age, constipation severity and physical inactivity, i.e. fewer hours of weekly sport (Fig 2A), were associated with an increase, and occupational solvent exposure with a decrease in α-diversity. Moreover, intake of thyroid medication, urate-lowering medication, dairy consumption and exhaustion from walking stairs (Fig 2B) were associated with a decreased α-diversity. None of the selected variables showed an interaction (p>.1) with physical inactivity on alpha diversity. When entering motor deficits into the regression, no association with alpha diversity was observed (p>.1); however, the interaction of motor deficits and physical inactivity showed a statistical trend (p=.088; Fig 2C).

**FIGURE 2.**
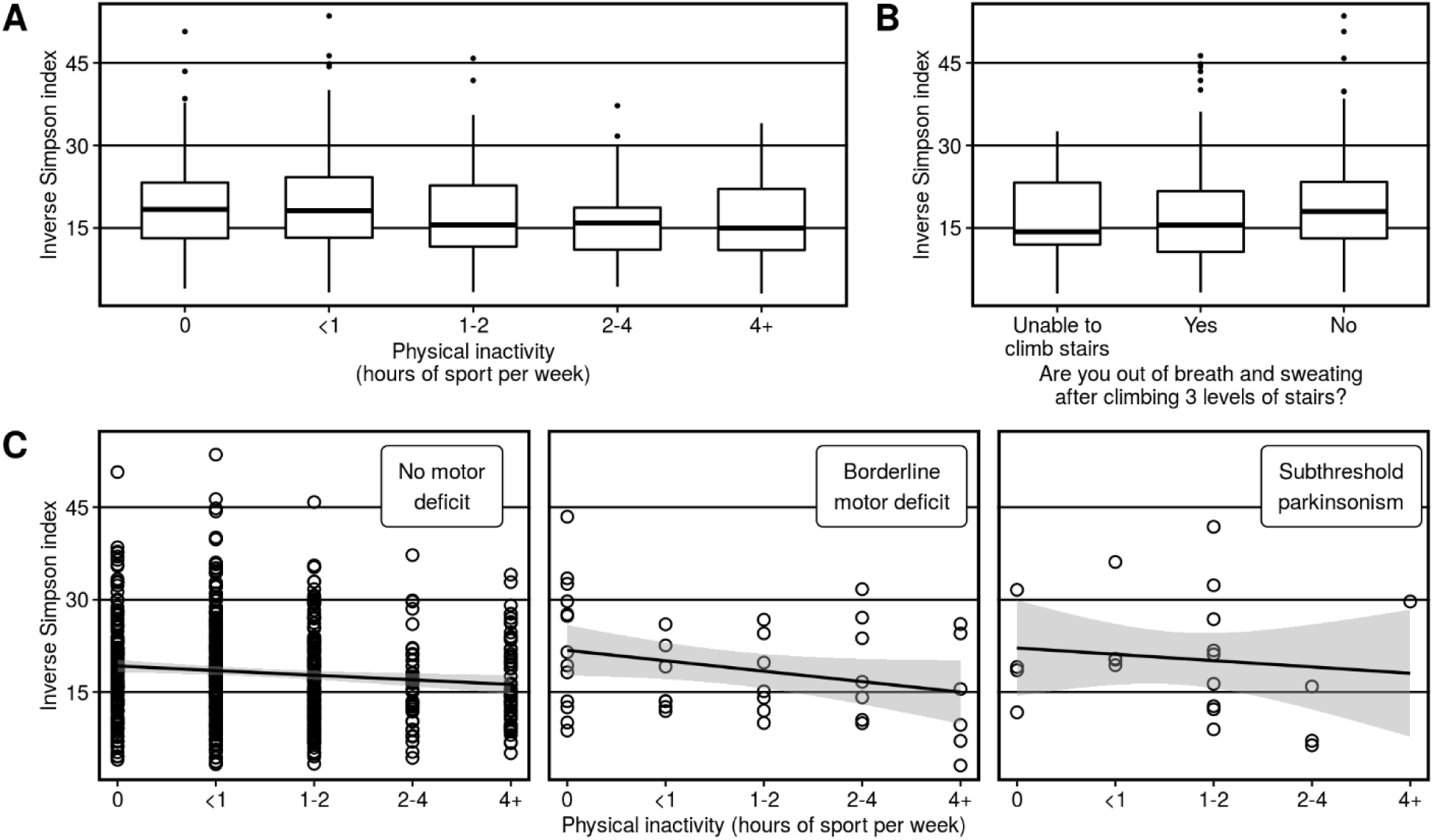
α-diversity and physical motor measures. Magnitude of α-diversity by (A) levels of weekly physical activity, (B) physical exhaustion, and (C) by physical activity levels in different motor deficit groups.

### β-diversity

Significant inter-sample differences in microbial composition as indicated by β-diversity were associated with age (Fig 3A) as well as multiple risk and prodromal markers of PD (Table 2). Among these markers, physical inactivity (Fig 3B) and constipation (Fig 3C) explained most of the variance (R²). Moreover, BMI, different medications, and dark bread consumption showed associations with β-diversity. While motor deficits showed no effect, the interaction between physical inactivity and motor deficits significantly explained variance in β-diversity.

**TABLE 2.** Associations of β-diversity with risk and prodromal markers in PD and additional factors.

|  | R<sup>2</sup> | F-value | p-value |  |
| --- | --- | --- | --- | --- |
| Age [years] | 0.008 | 4.88 | 0.001 | *** |
| Risk markers |  |  |  |  |
| Physical inactivity [h sport/week] | 0.014 | 9.09 | 0.001 | *** |
| Male sex [no/yes] | 0.006 | 3.59 | 0.003 | ** |
| Smoking [pack-years] | 0.004 | 2.36 | 0.020 | * |
| Prodromal markers |  |  |  |  |
| Constipation (severity sum score) | 0.009 | 5.48 | 0.002 | ** |
| Possible RBD [no/yes] | 0.003 | 2.14 | 0.037 | * |
| Subthreshold parkinsonism | 0.002 | 1.45 | 0.159 |  |
| Physical inactivity * Subthreshold parkinsonism | 0.006 | 3.56 | 0.002 | ** |
| Additional factors |  |  |  |  |
| Diabetes medication [no, yes] | 0.004 | 2.66 | 0.011 | * |
| Urate-lowering medication [no, yes] | 0.004 | 2.43 | 0.016 | * |
| Beta-blocker medication [no, yes] | 0.003 | 1.91 | 0.060 | (*) |
| Dark bread [no, yes] | 0.003 | 2.19 | 0.034 | * |
| BMI [kg/m<sup>2</sup>] | 0.013 | 8.41 | 0.001 | *** |

BMI = body mass index; RBD = REM-sleep behavior disorder, PD = Parkinson's disease; $p < .1$ (\*); $p < .05$ (\*); $p < .01$ (\*\*); $p < .001$ (\*\*\*)

**FIGURE 3.**
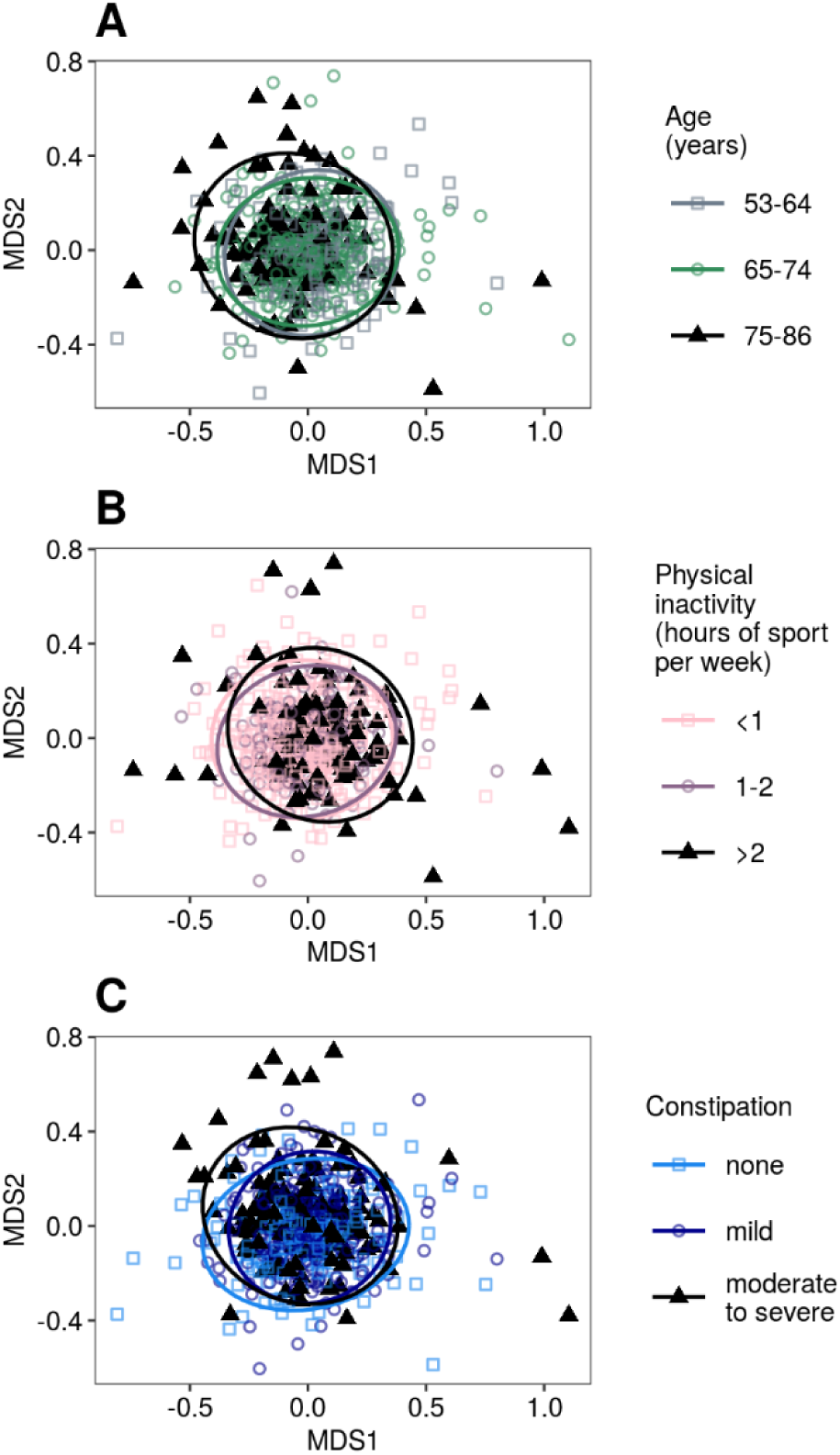
Non-metric multidimensional scaling (NMDS) plots in 2-dimensional space (MDS1 and 2) of factors showing significant Bray-Curtis distance differences. Differences in microbial composition between (A) age groups, (B) groups with different physical activity levels, and (C) groups stratified by severity of constipation are shown. Circles indicate 95% confidence intervals.

### Enterotypes

*Bacteroides*-enriched (70.9% of samples) microbiomes were more frequent compared to *Prevotella*-enriched (21.5%) and *Firmicutes*-enriched (7.7%) microbiomes. Enterotypes differed significantly in several risk and prodromal markers and additional factors (Table 3). For instance, the lowest physical activity levels as well as most severe constipation were observed for the *Firmicutes*-enriched enterotype. Significantly higher physical activity levels were observed in individuals with the *Bacteroides*-enriched compared to other enterotypes. Subthreshold parkinsonism was least frequently observed in individuals with the *Prevotella*-enriched enterotype, with a significant difference in *Prevotella*-*Firmicutes* enterotype comparisons (Fig 4).

**TABLE 3.**
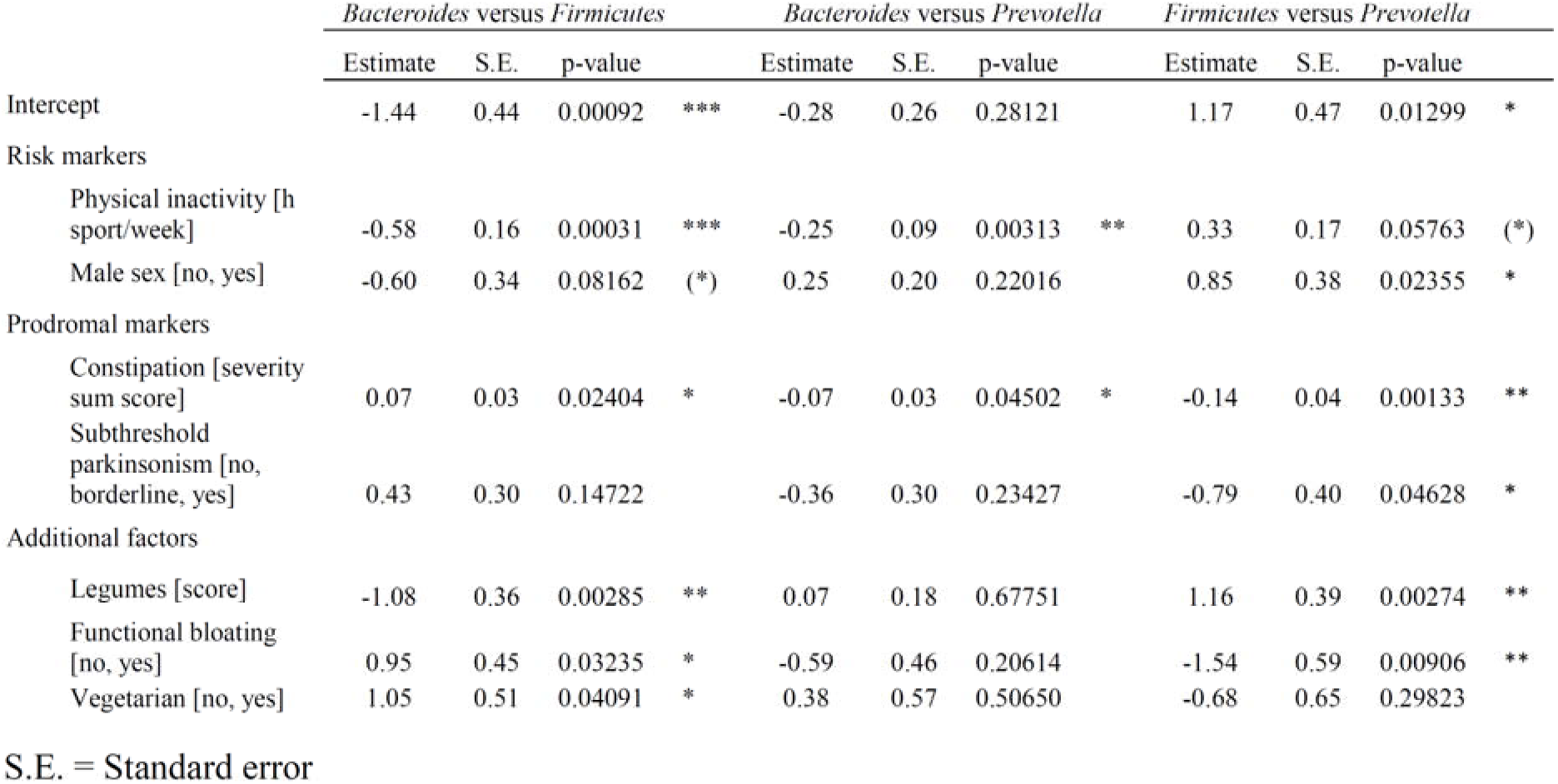
Associations of enterotypes with risk and prodromal markers of PD and additional factors.

**FIGURE 4.**
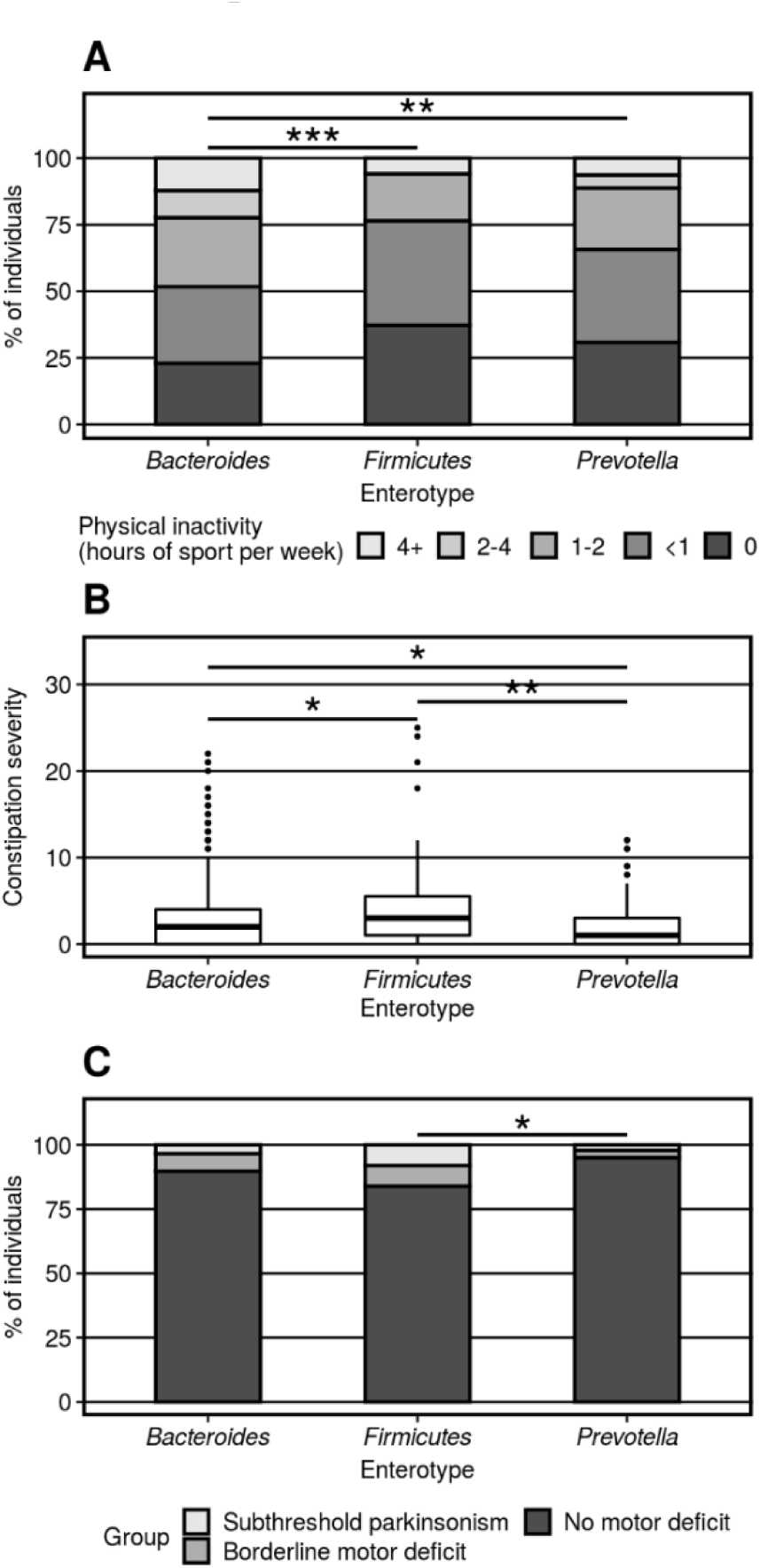
Enterotype group differences regarding (A) proportions of levels of physical activity, (B) severity of constipation, and (C) proportions of individuals with motor deficits indicating subthreshold parkinsonism.

### Differential abundance

Several of the variables associated with β-diversity were also linked to the abundances of specific taxa (Supporting materials). The variable with the longest list of differentially abundant taxa was severity of constipation (Table 4). Out of taxa previously associated with PD on the genus level^10^, constipation severity was significantly associated with decreased abundance of *Faecalibacterium* (adjusted p = .022), and physical exhaustion with a decrease in *Bifidobacterium* (adjusted p = .039). Further, motor deficits indicating subthreshold parkinsonism were associated with a decrease in *Odoribacter* (adjusted p = .031). Possible RBD was associated with a decrease in *Faecalicoccus*, *Lactobacillus* and *Victivallis*, and an increase in the abundance of *Haemophilus*. The taxa *Prevotella and Akkermansia* were not statistically significant in any of the differential abundance comparisons.

**TABLE 4.**
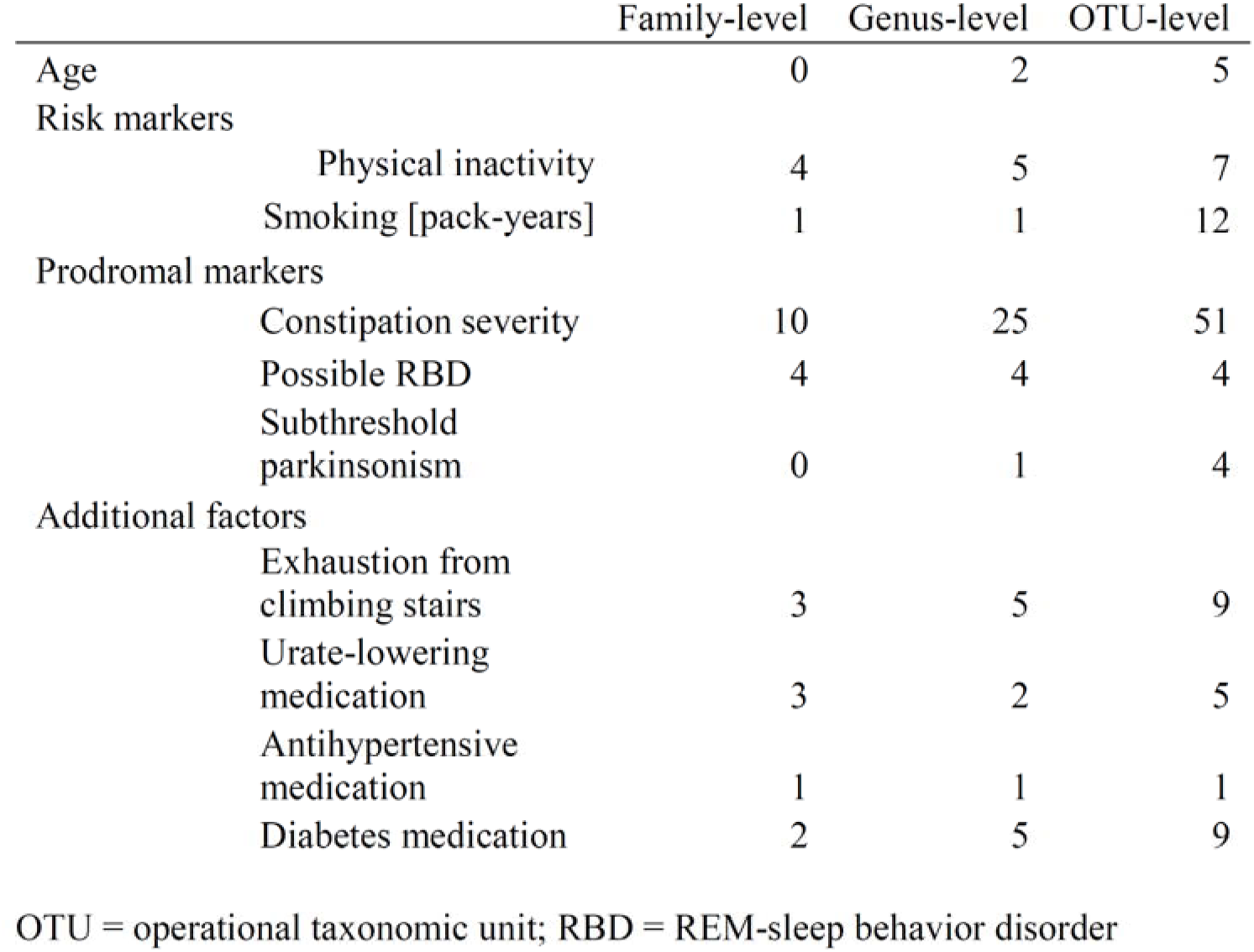
Number of differentially abundant taxa in association with risk and prodromal markers of PD and additional factors.

|  | Family-level | Genus-level | OTU-level |
| --- | --- | --- | --- |
| Age | 0 | 2 | 5 |
| Risk markers |  |  |  |
| Physical inactivity | 4 | 5 | 7 |
| Smoking [pack-years] | 1 | 1 | 12 |
| Prodromal markers |  |  |  |
| Constipation severity | 10 | 25 | 51 |
| Possible RBD | 4 | 4 | 4 |
| Subthreshold parkinsonism | 0 | 1 | 4 |
| Additional factors |  |  |  |
| Exhaustion from climbing stairs | 3 | 5 | 9 |
| Urate-lowering medication | 3 | 2 | 5 |
| Antihypertensive medication | 1 | 1 | 1 |
| Diabetes medication | 2 | 5 | 9 |

OTU = operational taxonomic unit; RBD = REM-sleep behavior disorder

## DISCUSSION

The present study investigated associations between gut microbial composition, risk markers and prodromal markers of PD, subthreshold parkinsonism as well as a wide range of potential confounders in 666 elderly individuals. Among these markers, particularly physical inactivity, constipation, possible RBD, and subthreshold parkinsonism were associated with significant alterations in microbial community composition. Moreover, age, male sex, smoking, and interactions of subthreshold parkinsonism and physical inactivity, showed significant associations with different microbial measures. As expected, effect sizes of individual markers and factors (≤1% explained variance) and multifactorial models (~8%) were small^16,17^. None of the microbial measures were associated with substantia nigra hyperechogenicity, olfactory loss, depression, orthostatic hypotension, urinary/erectile dysfunction, family history of PD or the overall prodromal PD probability calculated based on marker profiles of individuals.

The relative abundances of bacterial taxa were mostly similar to other samples of healthy elderly individuals^32^. Lower relative abundance of, e.g. *Bacteroides* compared to previous findings, may be explained by differences in cohort composition, e.g. the TREND cohort enrichment with individuals showing prodromal PD markers.

Age, constipation severity, and physical inactivity were associated with increasing within-sample α-diversity. Higher α-diversity in PD patients compared to controls has been reported^33–35^, yet overall evidence is inconsistent^10^ and it is unclear whether α-diversity and PD are linked. Similar to our results, older age in an elderly population has been associated with higher α-diversity^36^. However, statistical modeling of such effects is complicated by the diseases, medications and lifestyle changes often associated with advanced age. Regarding constipation severity, higher α-diversity has been linked to harder stool consistency^16^ and functional constipation^37^. Autonomous dysfunction related to prodromal PD may contribute to these α-diversity effects, which however need to be disentangled from other potentially important factors, such as diet, stool water content and transit time, and bacterial growth rates^38^. Physical inactivity increases the risk of many chronic diseases^20^ including PD^15^; conversely, being active may lead to lower prevalence of prodromal PD markers such as constipation^39^. The effects of inactivity could be related to the processes observed in PD, i.e. increased colonic inflammation^7^, immune responses^6^ and intestinal barrier permeability^5^. In our comparisons, the inactivity-associated increase in α-diversity was pronounced in individuals with borderline motor deficits and subthreshold parkinsonism. High α-diversity in elderly, inactive and constipated individuals could therefore be hypothesized as a microbial indicator of impending PD. Exhaustion from climbing stairs, an indicator of low cardiorespiratory fitness, was associated with a reduction in α-diversity, in line with earlier studies indicating lower diversity in lower fitness^40^ and frailty^41^. This apparently contradictory result may be partly explained by differences between physical activity and capability. Moreover, the PD risk marker occupational solvent exposure^15^ was associated with a decrease in α-diversity. Strengths and directions of effects of different risk and prodromal markers on microbial composition may vary based on the underlying pathophysiological pathways, potentially explaining why the overall prodromal PD probability calculated based on comprehensive marker profiles showed no significant association with α-diversity.

β-diversity (between samples) has been consistently shown to differ between PD patients and controls^10^. After initial exploratory screening analyses of 108 variables, age and several PD risk and prodromal markers remained significantly associated with β-diversity even in multifactorial models. These findings suggest that microbial composition might already be altered in the prodromal phase of PD. Although α- and β-diversity are not directly related, constipation and physical inactivity were again among the variables explaining most of the variance. As observed with α-diversity, physical inactivity and motor deficits indicating subthreshold parkinsonism showed an interaction effect. It remains speculative to which degree these associations play a specific role in prodromal PD. Possible links between β-diversity and RBD and the PD risk markers male sex and smoking further support the concept of microbial changes preceding PD. While replicating effects of antidiabetic and beta-blocker medication^17^, dark bread was the only dietary factor associated with β-diversity. Dark bread can be seen as an indicator of high fiber consumption, and thus presumably better intestinal barrier integrity and short-chain fatty acid (SCFA) production^42^. Since both have been suggested to be impaired in PD^5,43^ and to play a role for several processes along the microbiota-gut-brain axis^42^, dietary factors might thereby also be important in prodromal PD. After physical inactivity, BMI explained most variance in β-diversity. For some dietary factors linked to BMI, independent effects might have been too small to reach the threshold for statistical significance.

Subthreshold parkinsonism was least frequently present in individuals with the *Prevotella*-enriched compared to the Firmicutes-enriched enterotype. This finding is in line with the reduced abundance of *Prevotella* in PD^10^, in more severely progressing PD^11^, and in RBD^12^ compared to controls. Thus, the present study supports the relevance of *Prevotella* in prodromal PD. Microbial changes due to constipation are often argued to confound microbiome analysis of PD patients. The *Prevotella*-enriched enterotype was also the least common in individuals with high constipation severity scores. Subjects with high scores typically had the *Firmicutes*-enriched enterotype in accordance with previous findings^44^. Viewing constipation in PD as being linked to a disturbed enteric nervous system showing similar cellular changes as affected brain structures may suggest common relevance of *Prevotella* for prodromal dysautonomic and motor deficits. In this light, constipation may not be confounding, but reflecting a common pathogenic process. While constipation may exert effects on microbial composition via several plausible mechanisms, such effects may be different in diseased individuals as compared to healthy subjects, as recently demonstrated in PD for the association between α-diversity and stool consistency^11^. Physical inactivity was more frequently observed in individuals with the *Firmicutes*-enriched enterotype, whereas the *Bacteroides*-enriched enterotype was more frequent in active subjects. While consistent with some evidence gained from human^45^ and mouse studies^46^, processes underlying the links between physical activity, other prodromal markers in PD^39^, obesity, nutrition and microbial composition are complex and need to be further investigated.

The results of differential abundance analyses were less consistent with previous findings, as most taxa reported for PD^10^ and RBD^12^ showed no association with risk and prodromal markers in PD. In contrast with enterotype analyses, no significant differential abundance effect was observed for *Prevotella*, which may be partly explained by differences in covariates considered. Among other candidate genera, *Akkermansia* showed no significant association with any of the risk and prodromal markers. However, possible RBD showed several differentially abundant taxa, which have not been previously associated with PD^10^ or RBD^12^ (*Faecalicoccus, Victivallis* and *Haemophilus*). *Lactobacillus* was decreased in individuals with possible RBD, whereas an increase in PD patients has previously been shown repeatedly. It is possible that subtypes of prodromal PD exist with varying involvement of the gut (e.g., RBD representing a subtype with early autonomic denervation)^47^. Also, the microbiome may (differentially) change over time, i.e. from prodromal to clinically established PD. Such complexity may hamper the identification of (prodromal) PD-related microbiome signatures. However, these signatures may allow for early stratification of individuals based on their microbiome (and underlying or inherent pathologies), and early and targeted therapeutic interventions.

Constipation severity was significantly associated with decreased abundance of *Faecalibacterium*, a SCFA-producing taxon that can exert positive effects on the intestinal mucosa^48^ and is decreased in PD^10^. The finding of a decrease in *Odoribacter* in individuals with subthreshold parkinsonism has potential relevance for prodromal PD. *Odoribacter* is a taxon involved in SCFA production and tryptophan metabolism, and might be relevant for gastrointestinal integrity, and serotonergic bowel dysfunction (prolonged transit time) as well as central nervous dysfunction (anxiety) as suggested by findings in an autism mouse model^49^.

The present study has several limitations. (1) Some markers, and dietary and medication data were assessed using self-reports. While those assessments were structured and highly standardized, quantitative measures or medical records, such as weekly physical activity and dietary intakes, might be more accurate. (2) Interaction analyses were limited to those involving physical inactivity given its relevance as PD risk marker^15^ and for prodromal PD markers^39^. Further research on marker interactions and clusters is needed to better model the complexity of microbial associations. (3) Stool samples were not frozen immediately after defecation but after postal delivery, and the delay may constitute a technical confounder. However, collection tubes contained a DNA stabilizer and the impact of delayed freezing on microbial composition should therefore be minimal^11,50^.

In conclusion, several risk and prodromal markers in PD, in particular markers related to motor aspects, were associated with altered microbial α- and β-diversity, enterotypes, and bacterial abundance. Constipation, physical inactivity, possible RBD and subthreshold parkinsonism might be particularly relevant for the prodromal microbiome in PD. Yet, many other markers predictive of PD and overall prodromal PD probability values showed no significant association with any microbial measure. Prodromal microbial changes might only be observable in subgroups with specific marker constellations. The functional roles of these markers and associated microbiota for intestinal permeability, stool immune mediators, colonic inflammation, and the systemic interactions with the host organism need further investigation. The prodromal microbiome(s) in PD, temporal dynamics of microbiota towards PD diagnosis and etiological relevance in prodromal PD should be investigated based on prospective marker profiles and (multi)omics data in incident PD cases.

## Data availability

TREND data are available upon request. Microbiome sequence raw data is available under the ENA accession number PRJEB32920.

## Acknowledgements

We would like to thank the participants for their continued participation in the TREND study and providing their stool samples. We acknowledge the work of the numerous (doctoral) students and study nurses, who actively contributed to study organization, and data collection, entry and monitoring. The TREND organization team consists of Corina Maetzler, Susanne Nußbaum, Dr. Anna-Katharina von Thaler, Ulrike Sunkel, Christian Mychajliw and Ramona Täglich. We would also like to thank Albina Gashi, Ella Mustanoja, and Paula Collin for their work on biological sample processing at the DNA Sequencing Lab. Samples were obtained from the Neuro-Biobank of the University of Tuebingen, Germany (https://www.hih-tuebingen.de/en/about-us/core-facilities/biobank/). This biobank is supported by the local university, the Hertie Institute for Clinical Brain Research and the DZNE.

## Authors’ contributions

SH: Study design, programming, statistical analyses, interpretation of results, drafting of manuscript.

VA: Bioinformatics pipeline, figures, statistical & programming support, interpretation of results, revision of manuscript.

US: TREND study organization and database, revision of manuscript.

A-KvT: TREND study organization and database, revision of manuscript.

LP: Training of technicians, organisation and coordination of DNA-extraction, DNA-libraries and DNA sequencing pipelines, development of DNA barcoding approach, revision of manuscript.

SHa: Conception of dietary factor assessment, revision of manuscript.

KB: TREND study conception and supervision, revision of manuscript.

GWE: TREND study conception and supervision, revision of manuscript.

WM: TREND study conception and supervision, revision of manuscript.

DB: TREND study conception and supervision, revision of manuscript.

PA: Study conception, interpretation of results, revision of manuscript.

FS: Study conception, interpretation of results, revision of manuscript.

